# Precise oculocentric mapping of TMS-evoked phosphenes

**DOI:** 10.1101/2020.11.27.401828

**Authors:** Andrew E. Silva, Katelyn Tsang, Syeda Javeria Hasan, Benjamin Thompson

**Author notes:** Corresponding Author: Andrew E. Silva.

## Abstract

Transcranial Magnetic Stimulation (TMS) – evoked phosphenes are oculocentric; their perceived location depends upon eye position. We investigated the accuracy and precision of TMS-evoked phosphene oculocentric mapping and demonstrate that perceived phosphene locations map veridically to eye position, although there are considerable individual differences in the reliability of this mapping. Our results emphasize the need to carefully control eye movements when carrying out phosphene localization studies and suggest that individual differences in the reliability of the reported position of individual phosphenes must be considered.

## Introduction

Transcranial Magnetic Stimulation (TMS) is a non-invasive neurostimulation technique that involves magnetic induction of an electrical current within a relatively localized area of superficial neural tissue. When an individual pulse, or a train of individual pulses, is applied to the visual cortex, an illusory light percept known as a phosphene is often experienced (Meador et al., 1997). While there may be some systematic variation in the subjective appearance of evoked phosphenes in different early visual areas, the phosphene threshold, or the minimum stimulation output power required to evoke a phosphene response, is similar across early visual cortex (Salminen-Vaparanta et al., 2014). Stimulating early cortical areas can elicit phosphenes somewhat reliably, but it is more difficult to elicit phosphenes from later visual areas, such as V5 and LOC (Schaeffner & Welchman, 2017).

While it is vital to carefully account for the reliability and limitations inherent in the subjective reporting of phosphenes (Mazzi et al., 2017), phosphene thresholds and subjective judgements about phosphene characteristics have proven to be useful probes into the function and the excitability of visual cortex. Blind participants were demonstrated to perceive phosphenes, though at a reduced rate relative to normal control participants, depending on the level of function of the primary visual cortex (Cowey & Walsh, 2000; Gothe et al., 2002). Additionally, reduced phosphene thresholds were observed in normal-vision participants after short-term light deprivation, indicating increased cortical excitability (Boroojerdi, 2000). Cueing spatial attention toward the anticipate location of the phosphene was similarly suggested to increase cortical excitability (Bestmann et al., 2007).

Phosphenes evoked through direct, invasive cortical stimulation are oculocentric, moving with self-generated eye movements (Brindley & Lewin, 1968). This is perhaps expected, as primary visual cortex is a retinotopically-defined area with adjacent cortical areas encoding adjacent retinal locations. As a result, researchers have been continuously developing visual prostheses that employ electrical cortical stimulation to evoke phosphenes, providing blind individuals with some baseline level of visual information (Dobelle & Mladejovsky, 1974; Lewis et al., 2015; Ong & da Cruz, 2012; Tehovnik et al., 2005). These visual prostheses allow recovery of some light discrimination, but spatial and temporal information remains course (Niketeghad & Pouratian, 2019). Similarly, non-invasive TMS also evokes oculocentric phosphenes (Meyer et al., 1991), though TMS must penetrate the scalp and skull and may therefore elicit a noisier oculocentric mapping.

Studies involving TMS-evoked phosphenes often require that participants fixate a point, taking for granted that participants comply with fixation and that all evoked phosphenes will be positioned identically relative to fixation if TMS-coil position remains constant. However, the precision of a phosphene’s location with repeated trials, and the precision of the oculocentric mapping of TMS-evoked phosphenes, has not been thoroughly investigated. In the current study, we carry out a systematic investigation of the relationship between eye position and TMS-evoked phosphenes. Participants were directed to fixate individual points arranged in a grid while TMS was delivered to a fixed scalp location to induce phosphenes. Phosphene location was reported for each fixation point. We found that overall, TMS-evoked phosphenes mapped accurately to the point of fixation, though inter-participant variability was observed that could not be explained by eye movements or TMS-coil position alone.

## Method

### Participants

Twenty-one participants with self-reported normal vision were recruited and tested for reliable phosphenes and stable eye tracking. Nine participants (6F, 3M, ages: 21-41, median: 24) completed the study. Informed consent was obtained from all recruited participants, and all participants were treated in accordance with the Code of Ethics of the World Medical Association (Declaration of Helsinki). Participants received CAN$30.

### Transcranial Magnetic Stimulation

Biphasic Triple-pulse TMS with an inter-pulse interval of 100 ms was delivered using a MagPro X100 (MagVenture Farum, Denmark) stimulator with the MCF-B65 coil guided by the Brainsight frameless stereotaxic neuronavigation system (Rogue Research, Inc., Montreal, Canada). Coil placement errors were carefully monitored during every stimulation trial. Trials were repeated if the position error exceeded 2 mm or if the angle or twist error was greater than 5°. In total, 11 trials across 9 participants were repeated due to poor coil placement.

### Procedure

#### Phosphene Thresholding

Individual participants’ stimulation intensity was determined using a phosphene thresholding procedure to find the minimum intensity for which triple-pulse TMS evoked a phosphene five consecutive times. Participants were dark adapted for 20 minutes with eyes open and all room lights off. They were then instructed to fixate a dim point (2.50 cd/m^2^) on a solid dark background (1.75 cd/m^2^) presented on computer monitor with all other lights turned off. The TMS coil was placed 2 cm above the inion and the participant was stimulated with triple-pulse TMS at 40% maximum stimulator output, after which participants indicated whether they perceived a phosphene. If no phosphene was perceived after 2 attempts, the intensity was increased in 10% steps, to a maximum of 80%, and if the initial position failed to evoke any phosphenes, the coil was moved in increments of 2 cm, gradually testing points further from the original stimulation point. After a phosphene was perceived, participants used a computer mouse to indicate the location of the phosphene while maintaining fixation. the stimulation point was adjusted until the reported phosphene was no more than 2.62° (100 pixels) away from fixation. The intensity of the stimulation was then changed in increments of 5% until the minimum intensity required for the participant to report seeing 5/5 phosphenes was found. The resulting coil position and stimulation intensity were saved and used in the main experiment.

#### Behavioral Apparatus, Task, and Stimuli

The stimulus software was programmed using the Python package Psychopy (Peirce, 2007, 2009). Stimuli were presented on a 70 inch computer monitor with 1920 × 1080 pixel resolution. Participants were seated and used a chin and headrest for head stabilization. The viewing distance was 67 cm, and each pixel subtended roughly 0.027 degrees.

To begin each trial, participants fixated a dim gray dot. Possible locations for fixation were defined using a 38.5 degree by 19.2 degree grid of points, with 7 equally-spaced points distributed horizontally and 5 equally-spaced points distributed vertically. Therefore, 35 total fixation locations were tested. On any given stimulation trial, only one randomly sampled fixation dot was visible. After TMS stimulation, participants used the mouse to indicate the location of the phosphene while maintaining fixation. If no phosphene was perceived, the stimulation was repeated before the mouse click. The stimulus intensity was increased by 2% of the maximum stimulator output if a phosphene was not evoked after two concurrent attempts during the experimental task

Participants always fixated the middle location during the first five stimulation trials. After these initial trials, all fixation locations, including the middle location, were tested three times. In all, 35 (fixation locations) × 3 (# trials) + 5 (initial middle trials) = 110 trials were presented. A Gazepoint GP3 eyetracker was used to monitor eye movements, and any trials exhibiting a saccade, defined as an eye movement with velocity greater than 40 degrees/s, were removed from analysis.

Before performing the main stimulation task, participants underwent a brief training procedure without TMS-evoked phosphenes. Instead, brief dim circular flashes (2.5 cm/m^2^) were presented to participants within 200 pixels or 5.3 degrees of fixation, and participants were required to indicate the location of the phosphene with a mouse click while maintaining fixation. The fixation point was randomly selected as in the main task.

## Results and Discussion

During the main experiment, 52 total stimulation trials failed to evoke a phosphene out of 110 (trials) × 9 (participants) = 990 total trials. Of these, 29 trials originated from participant A5 (See Figure 1). All failed trials were repeated successfully, and all participants reported seeing the 110 phosphenes required to complete the study. Figures 1A and 1B present single-subject and group results, respectively. Before any processing, all data were centered relative to the arithmetic mean of all within-subject absolute distances between the middle fixation location and the phosphenes elicited when centrally fixating. In other words, for each subject, the grid was aligned to the average position of the phosphene corresponding to the central fixation point. Figure 1A presents the reported individual phosphene locations for each participant. Figure 1B presents the arithmetic mean of each participant’s phosphene reports at each fixation location.

**Figure 1:**
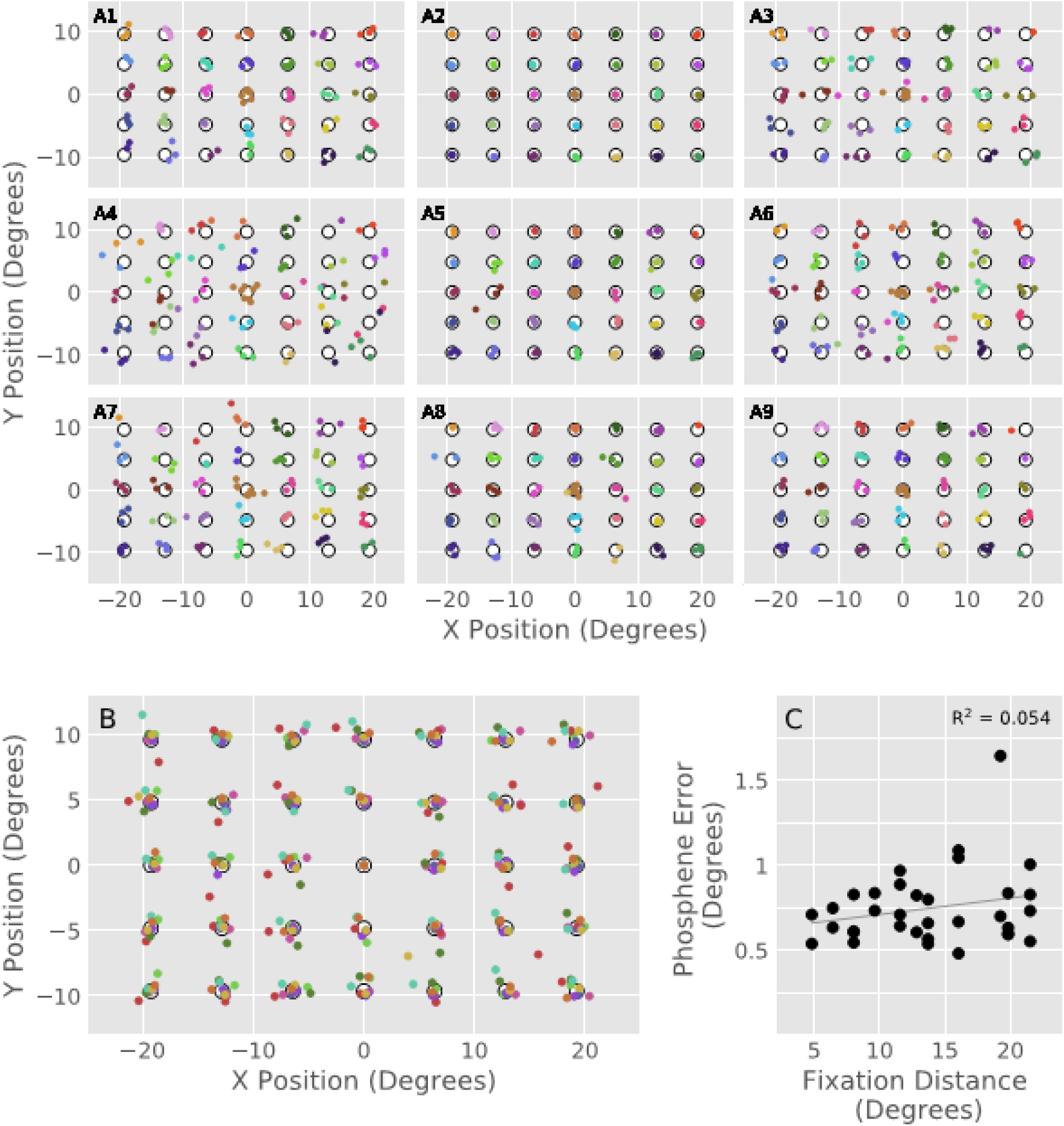
Oculocentric phosphene mapping results. All phosphene reports were centered relative to the arithmetic mean of the individual within-subject phosphene locations at the center-most fixation location. A) Individual-subject results. Identically colored dots represent individual phosphene reports evoked when fixating the same location. B) Average reported phosphene locations at each oculocentric position, calculated as the arithmetic mean of all evoked phosphenes at the given position per participant. Identically colored dots represent data from the same participant. C) Scatterplot illustrating the overall relationship between phosphene position error and eccentric distance of the given oculocentric position to the middle fixation location. No significant association was found.

Inspection of Figures 1A and 1B reveals relatively accurate oculocentric mapping of perceived phosphene location to fixation position. To test whether phosphene position relative to fixation varied as a function of fixation eccentricity, a phosphene error was defined as the average absolute distance between fixation and phosphene location for each fixation location across participants. A linear regression was carried out on the phosphene error data against the distance of the associated fixation location from the central position. No significant relationship between phosphene error and fixation position was found, R^2^ = 0.05, *p* = 0.19; Figure 1C illustrates this analysis.

Visual inspection of Figure 1A demonstrates that the variability of the location of elicited phosphenes varied noticeably between participants. Potential explanations for variable phosphene scatter include external factors, such as TMS coil position error and incorrect eye movements. The current study controlled for these factors by monitoring TMS coil position and removing trials with saccades. Even so, participants A3, A4, A6, and A7 exhibited noticeably more scatter in their phosphene reports than the other participants. To investigate whether systematic differences exist between our high and low scatter participants with respect to the oculocentric mapping of phosphenes, two post-hoc linear regressions using phosphene errors calculated with data from only the high-scatter participants and only the low-scatter participants were carried out identically to the group analysis, neither finding a significant effect, R^2^ = 0.02, *p* = 0.46, and R^2^ = 0.08, *p* = 0.11, respectively.

These results confirm that on average, TMS-evoked phosphenes are oculocentric, varying their position tightly with respect to eye movements (Brindley & Lewin, 1968; Meyer et al., 1991). No systematic changes in phosphene position accuracy were found as a function of the eccentricity of fixation. Furthermore, these results demonstrate the presence of individual differences with respect to the positional stability of TMS-evoked phosphenes, even when reasonable measures are taken to minimize obvious confounding factors, such as eye movements and TMS coil position errors. Ultimately, the results of our study reinforce the need to carefully control fixation in studies that involve TMS-evoked phosphene localization and to account for inter-subject differences in the reliability of phosphene position reports.

## Notes

### Competing Interest Statement

The authors have declared no competing interest.

## References

Bestmann, S., Ruff, C. C., Blakemore, C., Driver, J., & Thilo, K. V. (2007). Spatial Attention Changes Excitability of Human Visual Cortex to Direct Stimulation. Current Biology, 17(2), 134–139. https://doi.org/10.1016/j.cub.2006.11.063

Boroojerdi, B. (2000). Enhanced Excitability of the Human Visual Cortex Induced by Short-term Light Deprivation. Cerebral Cortex, 10(5), 529–534. https://doi.org/10.1093/cercor/10.5.529

Brindley, G. S., & Lewin, W. S. (1968). The sensations produced by electrical stimulation of the visual cortex. The Journal of Physiology, 196(2), 479–493. https://doi.org/10.1113/jphysiol.1968.sp008519

Cowey, A., & Walsh, V. (2000). Magnetically induced phosphenes in sighted, blind and blindsighted observers. In NeuroReport (Vol. 11, Issue 14, pp. 3269–3273). https://doi.org/10.1097/00001756-200009280-00044

Dobelle, W., & Mladejovsky, M. (1974). Phosphenes produced by electrical stimulation of human occipital cortex, and their application to the development of a prosthesis for the blind. Journal of Physiology, 243, 553–576. https://doi.org/jphysiol.1974.sp010766

Gothe, J., Brandt, S. A., Irlbacher, K., Röricht, S., Sabel, B. A., & Meyer, B. U. (2002). Changes in visual cortex excitability in blind subjects as demonstrated by transcranial magnetic stimulation. In Brain (Vol. 125, Issue 3, pp. 479–490). https://doi.org/10.1093/brain/awf045

Lewis, P. M., Ackland, H. M., Lowery, A. J., & Rosenfeld, J. V. (2015). Restoration of vision in blind individuals using bionic devices: A review with a focus on cortical visual prostheses. Brain Research, 1595, 51–73. https://doi.org/10.1016/j.brainres.2014.11.020

Mazzi, C., Savazzi, S., Abrahamyan, A., & Ruzzoli, M. (2017). Reliability of TMS phosphene threshold estimation: Toward a standardized protocol. Brain Stimulation, 10(3), 609–617. https://doi.org/10.1016/j.brs.2017.01.582

Meador, K. J., Ray, P. G., & Loring, D. W. (1997). Physiology of perception: parameters of TMS-induced phosphenes. Electroencephalography and Clinical Neurophysiology, 102(1), P12. https://doi.org/10.1016/S0013-4694(97)86260-8

Meyer, B. U., Diehl, R., Steinmetz, H., Britton, T. C., & Benecke, R. (1991). Magnetic stimuli applied over motor and visual cortex: influence of coil position and field polarity on motor responses, phosphenes, and eye movements. Electroencephalography and Clinical Neurophysiology. Supplement, 43, 121–134. http://www.ncbi.nlm.nih.gov/pubmed/1773752

Niketeghad, S., & Pouratian, N. (2019). Brain Machine Interfaces for Vision Restoration: The Current State of Cortical Visual Prosthetics. Neurotherapeutics, 16(1), 134–143. https://doi.org/10.1007/s13311-018-0660-1

Ong, J. M., & da Cruz, L. (2012). The bionic eye: A review. Clinical and Experimental Ophthalmology, 40(1), 6–17. https://doi.org/10.1111/j.1442-9071.2011.02590.x

Peirce, J. W. (2007). PsychoPy-Psychophysics software in Python. Journal of Neuroscience Methods, 162, 8–13. https://doi.org/10.1016/j.jneumeth.2006.11.017

Peirce, J. W. (2009). Generating Stimuli for Neuroscience Using PsychoPy. Frontiers in Neuroinformatics, 2, 10. https://doi.org/10.3389/neuro.11.010.2008

Salminen-Vaparanta, N., Vanni, S., Noreika, V., Valiulis, V., Móró, L., & Revonsuo, A. (2014). Subjective characteristics of TMS-induced phosphenes originating in human V1 and V2. Cerebral Cortex, 24(10), 2751–2760. https://doi.org/10.1093/cercor/bht131

Schaeffner, L. F., & Welchman, A. E. (2017). Mapping the visual brain areas susceptible to phosphene induction through brain stimulation. In Experimental Brain Research (Vol. 235, Issue 1, pp. 205–217). https://doi.org/10.1007/s00221-016-4784-4

Tehovnik, E. J., Slocum, W. M., Carvey, C. E., & Schiller, P. H. (2005). Phosphene induction and the generation of saccadic eye movements by striate cortex. Journal of Neurophysiology, 93(1), 1–19. https://doi.org/10.1152/jn.00736.2004

